# Vegetable oil surfactants are synergists that can bias neonicotinoid susceptibility testing in adult mosquitoes

**DOI:** 10.1101/2023.04.18.537421

**Authors:** Fred A. Ashu, Caroline Fouet, Marilene M. Ambadiang, Véronique Penlap-Beng, Colince Kamdem

## Abstract

**Background:** The standard operating procedure for testing the susceptibility of adult mosquitoes to clothianidin, a neonicotinoid, recommends using a vegetable oil ester as surfactant. However, it has not yet been determined if the surfactant is an inert ingredient or if it can act as a synergist and bias the test.

**Methodology/Principal Findings:** Using standard bioassays, we tested the synergistic effects of a vegetable oil surfactant on a spectrum of active ingredients including four neonicotinoids (acetamiprid, clothianidin, imidacloprid and thiamethoxam) and two pyrethroids (permethrin and deltamethrin). Three different formulations of linseed oil soap used as surfactant were far more effective than the standard insecticide synergist piperonyl butoxide in enhancing neonicotinoid activity in *Anopheles* mosquitoes. At the concentration used in the standard operating procedure (1% v/v), vegetable oil surfactants lead to more than 10-fold reduction in lethal concentrations, LC_50_ and LC_99_, of clothianidin in a multi-resistant field population and in a susceptible strain of *Anopheles gambiae*. At 1% or 0.5% (v/v), the surfactant restored susceptibility to clothianidin, thiamethoxam and imidacloprid and increased mortality to acetamiprid from 43 ± 5.63% to 89 ± 3.25% (P<0.05) in resistant mosquitoes. By contrast, linseed oil soap had no effect on the level of resistance to permethrin and deltamethrin suggesting that the synergism of vegetable oil surfactants may be specific to neoniconoids.

**Conclusions/Significance:** Our findings indicate that vegetable oil surfactants are not inert ingredients in neonicotinoid formulations, and their synergistic effects undermine the ability of standard testing procedures to detect early stages of resistance.

## 1. Introduction

The development of new chemicals is crucial to preserve the efficacy of proven malaria vector control tools such as long-lasting insecticidal nets and indoor residual spraying [1,2]. As new products are identified and tested, evaluating their efficacy in wild vector populations is of prime importance. This process involves using reference susceptible strains to determine discriminating doses of the insecticide that will be tested on wild mosquito populations to assess their susceptibility profiles [3,4].

Neonicotinoids provide promising alternatives to control mosquito populations that have developed resistance to insecticides previously deployed in large-scale malaria control efforts [5,6]. Two formulations containing clothianidin, a neonicotinoid, have been pre-qualified for indoor residual spraying by the World Health Organization (WHO) [7]. These formulations are SumiShield® (Sumitomo Chemical Company, Japan) and Fludora® Fusion (Bayer Crop Science, Monheim, Germany), a combination of clothianidin and deltamethrin [8,9]. Two standard testing procedures are currently available for monitoring susceptibility in wild anopheline populations: a WHO paper-test, which assesses mortality in young adult females exposed to SumiShield® and a bottle bioassay which evaluates their tolerance to clothianidin dissolved in a solvent [3,4,10].

Early efforts to implement a standard testing procedure were challenged by the low solubility of clothianidin observed when large quantities of active ingredient were being diluted in water, ethanol or acetone [10,11]. To circumvent this issue, it has become common to add a surfactant to insecticide solutions used for bottle bioassays [2,12–14]. The WHO standard operating procedure for testing clothianidin susceptibility of adult mosquitoes recommends mixing the active ingredient with 1% (v/v) of a formulation of rapeseed fatty acid esters (Mero®, Bayer, Reading, UK) to improve solubility and uptake of the insecticide by mosquitoes [4]. Surfactants act as surface-active agents and have been widely used in agricultural pest management to solubilize, suspend, or disperse active ingredients allowing more pesticide to reach its target in aqueous solutions [15,16]. Surfactants and other adjuvants used in pesticide formulations are often referred to as “inert” ingredients, but some can have biological activity on their own or can enhance the toxicity of insecticides, so that they jointly exert a larger effect than predicted—a phenomenon called synergy [16].

The purpose of susceptibility testing is to gain information on the ability of wild populations to adapt to selection pressure generated by insecticide treatments. For new insecticides such as clothianidin, early detection of reduction in susceptibility, even at low levels, is crucial to guide monitoring efforts that would help prevent the emergence and spread of resistance [17]. If the surfactant used in standard testing procedures is a synergist of clothianidin, the mixture may become extremely potent thereby reducing the test’s ability to detect early stages of resistance. Thus, it is necessary to have a better understanding of the spectrum of interactions between the surfactant and the active ingredient as well as the confounding effects these interactions may have on susceptibility evaluation. The aim of the present study was to investigate the effects of a vegetable oil surfactant on the potency of clothianidin, acetamiprid, imidacloprid, thiamethoxam, permethrin and deltamethrin. We compared mortality rates induced by the active ingredient alone or by the insecticide-surfactant mixture in a multiresistant field population of *Anopheles gambiae* whose larvae and adults have developed resistance to neonicotinoids [18–20]. We found that vegetable oil surfactants have little effect on resistance to pyrethroids, but are very effective synergist of neonicotinoids.

## 2. Methods

### 2.1 Mosquito populations

The study focused on *An. gambiae* mosquitoes collected from Nkolondom (3°56’43” N, 11°31’01” E) in the suburban area of Yaoundé, Cameroon. Approval to conduct a study in the Center region (N°: 1-140/L/MINSANTE/SG/RDPH-Ce), ethical clearance (N°: 1-141/CRERSH/2020) and research permit (N°: 000133/MINRESI/B00/C00/C10/C13) were granted by the ministry of public health and the ministry of scientific research and innovation of Cameroon. Due primarily to chronic exposure to agricultural pesticides, larvae and adult mosquitoes from Nkolondom are resistant to neonicotinoids, pyrethroids and several organophosphates and carbamates [18–24]. Immature stages of this super-resistant population were collected from standing water in furrows between ridges across a large farm using a dipper [25]. *An. gambiae* larvae were transported in plastic containers to the insectary where they were reared under standard environmental conditions (25 ± 2°C, 70-80% relative humidity, light: dark cycles of 12:00 h each) while being fed with TetraMin^®^. Adults that emerged were maintained in 30×30×30 cm Bug Dom cages and were provided with 10% sugar solution. Female adults aged between 2 and 3 days were used for insecticide resistance bioassays. The susceptible laboratory strain *An. gambiae* Kisumu, reared in the insectary under standard conditions was used for comparison.

### 2.2 Insecticides and surfactants

In this study, we used technical grade of four neonicotinoids (clothianidin, acetamiprid, imidacloprid and thiamethoxam) and two pyrethroids (permethrin and deltamethrin) all obtained from Sigma-Aldrich (Pestanal®, Sigma-Aldrich, Dorset, United Kingdom). We chose six insecticides for which we had sufficient information on the level of resistance in *Anopheles* mosquitoes from Nkolondom. As *An. gambiae* larvae and adults from this agricultural site are highly resistant to the six active ingredients tested, this allowed us to assess synergism by detecting any increase in mortality due to the surfactant [18– 20,22,24,26]. The insecticides were diluted in absolute ethanol except imidacloprid, which was soluble in acetone. Three commercial formulations of linseed oil soap were purchased from supermarkets in Yaoundé and used as surfactant: (1) Maître Savon de Marseille®, Saint-Laurent-du-Var, France (S1), (2) Carolin Savon noir®, Bolton, Belgium (S2), (3) La perdrix®, Paris, France (S3). Insecticide solutions containing soap were prepared and stored at 4°C for at least 24 h before being used. A solution of ethanol or acetone containing soap was used as control in bioassay tests.

### 2.3 Bottle bioassays

We investigated the synergistic effects of soap using Center for Disease Control (CDC) bottle bioassays to compare mortality values of *An. gambiae* mosquitoes exposed either to the insecticide alone or to a mixture containing the active ingredient and 1% (v/v) linseed oil soap. Mortality would significantly increase in the presence of soap in case of synergism. Wheaton 250-ml bottle and its cap were coated with 1 ml of insecticide solution following the CDC protocol [27]. For each bioassay, we used four test bottles coated with test insecticide and two control bottles. The discriminating concentrations defined as the lowest dose of the insecticide required to kill 100% of exposed individuals from a susceptible population in 24 h were obtained from published protocols. The discriminating concentrations tested were: 12.5 μg/ml deltamethrin, 21.5 μg/ml permethrin, 75 μg/ml acetamiprid, 150 μg/ml clothianidin, 200 μg/ml imidacloprid and 250 μg/ml thiamethoxam [4,19,27,28]. Control solutions consisted of solvent (ethanol or acetone) or solvent containing 1% (v/v) soap.

To perform bioassay tests, 25 female mosquitoes, 3-5 day old, were aspired from cages and gently released into paper cups and then into the treated bottles for 1 h. The mosquitoes were then transferred into holding paper cups and provided with 10% sugar solution, and mortality was recorded after 24 h. We carried out different types of test to better understand the interactions between the active ingredient and soap: (1) a test to evaluate the effects of three different brands of linseed oil soap (1% v/v) on the susceptibility of the super-resistant *An. gambiae* population exposed to 150 µg/ml clothianidin, (2) a test comparing the effects of 1% (v/v) soap on resistance to four different neonicotinoids and two pyrethroids (3) an evaluation of the effect of halving the concentration of soap or pre-exposing mosquitoes to the surfactant (4) and finally an analysis of the effect of soap on the short-term lethal toxicity of clothianidin. In the last test, we subjected adults to gradual concentrations of clothianidin containing 1% soap and we compared the 24-h lethal concentrations (LC_50_ and LC_99_) between the resistant field population and the susceptible strain *An. gambiae* Kisumu. The doses tested were as follows: *An. gambiae* Kisumu (0.39, 0.78, 1.56, 3.12, 4.68, 6.25, 9.37, 12.5, 25 μg/ml); *An. gambiae* Nkolondom (1.56, 3.12, 4.68, 6.25, 12.5, 25, 100 μg/ml). Bioassays were performed under a controlled environment of 25–27°C, 70–90% relative humidity and a 12:12h light/dark photoperiod.

### 2.4 Data analysis

The mortality rate (%) was used to estimate the susceptibility of field-collected and laboratory mosquitoes. Mortality was averaged across four replicates for each insecticide dose and a log-logistic model was used to fit the dose-response curve with the *drc* package in R version 4.2.2 [29]. A probit model was implemented to calculate LC_50_ and LC_99_ with their 95% confidence intervals using the *ecotox* package in R. LC_99_ and LC_50_ corresponded to the dose of active ingredient at which respectively 99% and 50% of individuals died after 1 h exposure and 24 h holding period. This analysis aimed to test if the lethal toxicity of the insecticide-surfactant blend was different between the susceptible strain and the multi-resistant filed population. The lethal toxicity was considered different if there was no overlap between the corresponding 95% confidence intervals of LC_50_. We used Wilcoxon rank sum test to determine if there was a significant difference in mortality rates across tests. All analyses were performed using R V. 4.2.2 [29].

## 3. Results

### 3.1 Three different brands of linseed oil soap synergized clothianidin

In order to determine if vegetable oil surfactants had the power to improve the activity of clothianidin, we tested the potential of three different commercial brands of linseed oil soap. The results showed that at 1% (v/v) any of the three brands could enhance the efficacy of clothianidin at a discriminating dose of 150 μg/ml. The mortality induced in the multi-resistant *An. gambiae* population collected from Nkolondom increased from 30 ± 3.49% within 24 h of exposure to clothianidin alone to 100% when 1% soap was added to the active ingredient (Figure 1). This result indicated that linseed oil soap was not an inert ingredient and had synergistic interactions with clothianidin. This first test also showed that this synergism is reproducible with three different brands of linseed oil soap. Mortality varied between 0 and 4% in all the control tests confirming that soap *per se* did not have insecticidal activity against *Anopheles*.

**Figure 1:**
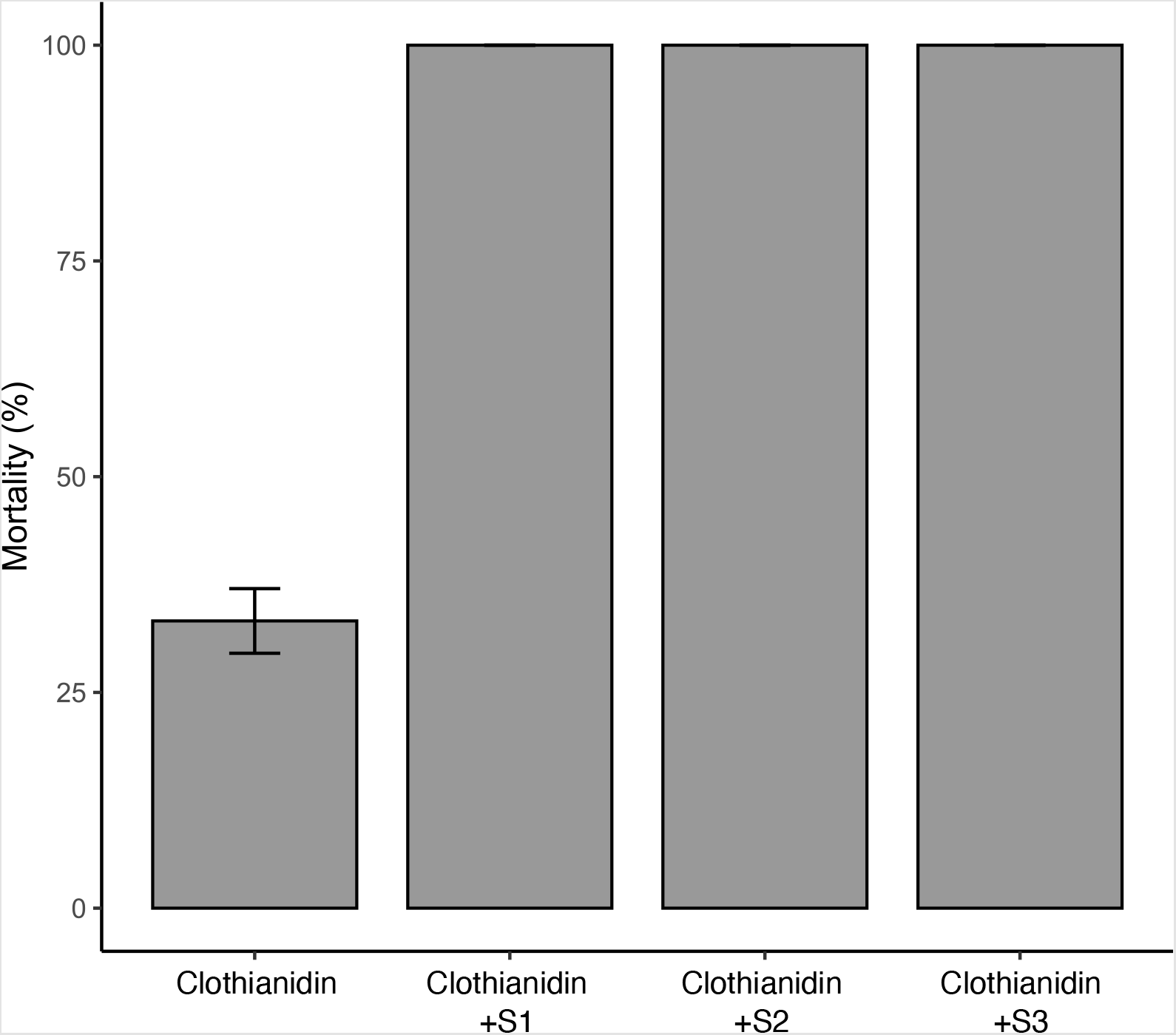
Mortality rate obtained within 24 h of exposure to clothianidin alone or to a mixture containing 150 µg/ml clothianidin and 1% (v/v) of one of the commercial formulations of linseed oil soap (S1, S2 and S3). Error bars represent the standard error of the mean.

### 3.2 Soap enhanced 24-h lethal toxicity of clothianidin

Focusing on clothianidin, we measured to what extent the surfactant enhanced the potency of neonicotinoids. To do so, we tested increasing concentrations of clothianidin containing 1% (v/v) soap (S1) and we used dose-response curves and a probit model to evaluate lethal toxicity in 24 h. As indicated by a previous study, when clothianidin is dissolved in ethanol without any surfactant, LC_99_ and LC_50_ against *An. gambiae* Kisumu are ∼143 μg/mL and 27 μg/ml, respectively [28]. In the present study, we found that there was a more than 10-fold reduction in the lethal concentrations when mixtures containing 1% soap were used (LC_99_: 19.70 μg/ml *CI*95%[11.20, 51.10] and LC_50_: 0.93 μg/ml [0.58, 1.30]) (Figure 2). In the presence of soap, short-term lethal toxicity of clothianidin also drastically increased in the multi-resistant Nkolondom population. Within 24 h of exposure, LC_99_ and LC_50_ against the resistant population were 13.10 μg/ml [7.95, 45.60] and 2.23 μg/ml [1.49, 2.86] respectively. Importantly, 95% confidence intervals of LC_99_ overlapped between *An. gambiae* Kisumu and field mosquitoes, which clearly indicated that susceptibility was restored in the presence of the surfactant. However, LC_50_ against the Nkolondom population was significantly higher compared to *An. gambiae* Kisumu as indicated by non-overlapping confidence intervals. This implied that 50% mortality was still more difficult to achieve in resistant mosquitoes even with the synergistic effect of soap.

**Figure 2:**
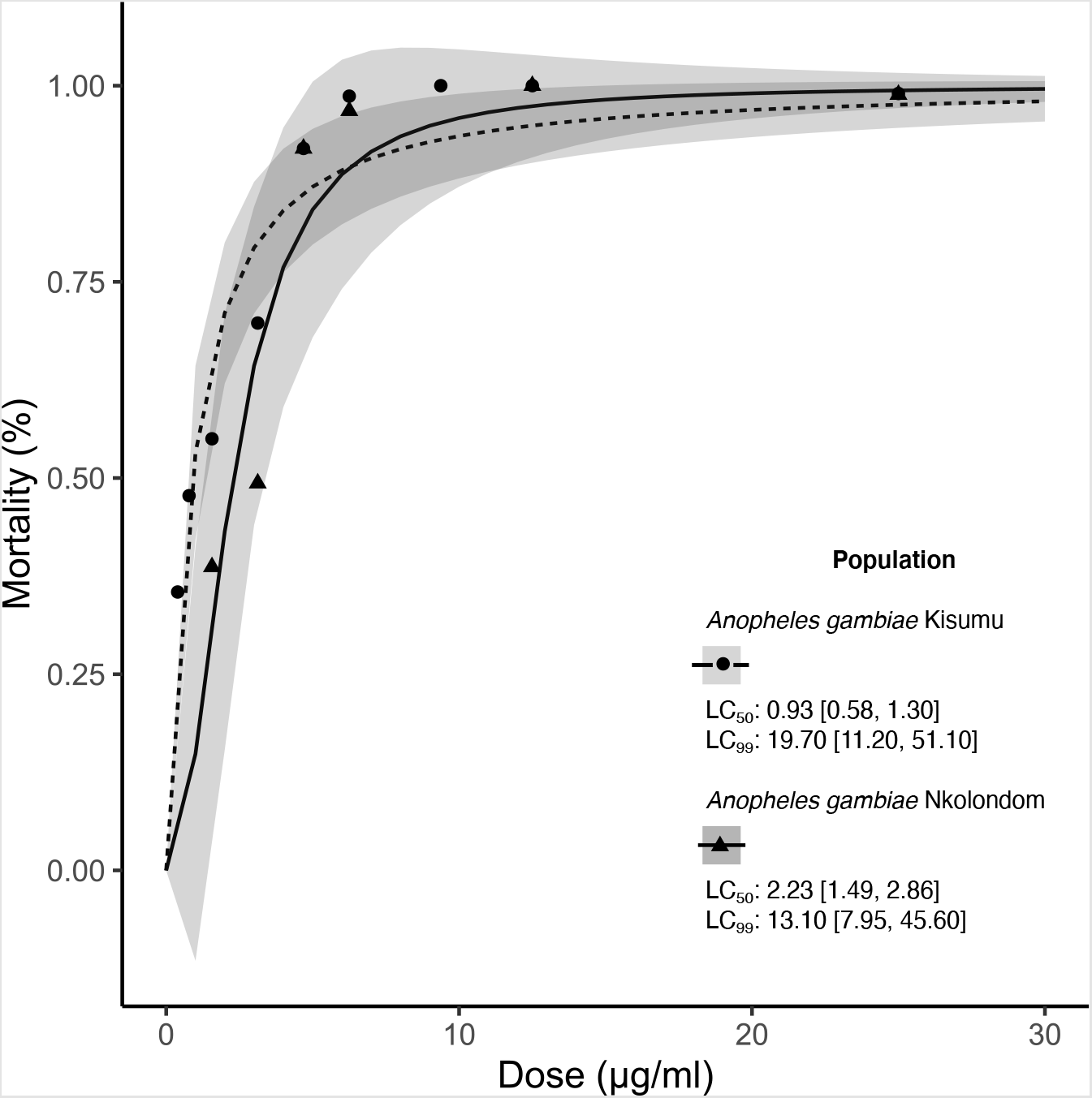
Dose-response curves with the standard error of the regression model (grey bands) describing the 24-h lethal toxicity of the mixture containing clothianidin (150 µg/ml) and 1% (v/v) surfactant. LC_50_ and LC_99_ were compared between the susceptible strain *An. gambiae* Kisumu and a multi-resistant population of *An. gambiae* collected from Nkolondom.

### 3.3 The surfactant restored susceptibility to neonicotinoids but not pyrethroids

Clothianidin is only moderately soluble in ethanol, even though the technical grade used was soluble when the mixture was allowed to rest for at least 24 h [18,19,28]. Therefore, since linseed oil soap reduces surface tension and increases the solubility of the active ingredient, it contributes to the release of more pesticide in the solution, which enhances activity. To test if reducing the concentration of surfactant affects synergism, we added 0.5% instead of 1% v/v soap to 8 μg/ml and 15 μg/ml of clothianidin and we measured mortality induced in resistant mosquitoes. Both concentrations were chosen within the 95% confidence intervals of LC_99_. 100% mortality was achieved within 24 h of exposure to each concentration, which showed that halving the dose of surfactant had no effect on synergism. In a separate experiment, we first exposed resistant mosquitoes for 1 h in bottles coated with 1% soap (S1) before releasing them for 1 h in bottles containing 8 μg/ml of clothianidin. Pre-exposure to soap did not significantly increase mortality (from 30 ± 3.49% to 55 ± 6.60%, Wilcoxon rank sum test, P>0.05), confirming that soap is more effective as a surfactant mixed with the neonicotinoid insecticide.

To determine if vegetable oil surfactants could induce broad-spectrum synergism beyond improving clothianidin solubility, we tested the effect of linseed oil soap on the activity of other insecticides with different chemical properties and solubility in aqueous solutions. We first measured the baseline susceptibility of the Nkolondom mosquito population to imidacloprid, acetamiprid, thiamethoxam, permethrin and deltamethrin. We then compared mortality rates between tests that used the active ingredient alone and bioassays involving a mixture of the insecticide and 1% (v/v) soap (S1). Figure 3 shows that susceptibility (100% mortality) to thiamethoxam and imidacloprid was restored within 24 h of exposure to the insecticide-soap mixture, while mortality to acetamiprid increased from 43 ± 5.63% to 89 ± 3.25% (Wilcoxon rank sum test, p<0.05). Meanwhile when linseed oil soap was added to deltamethrin or permethrin, a slight but not significant reduction in mortality was observed (Wilcoxon rank sum test, p>0.05) suggesting that there was no synergism between this surfactant and pyrethroids. Taken together, vegetable oil surfactants are broad-spectrum synergists which enhance the activity of neonicotinoids against *An. gambiae* mosquitoes, but have little effect on the efficacy of pyrethroids.

**Figure 3:**
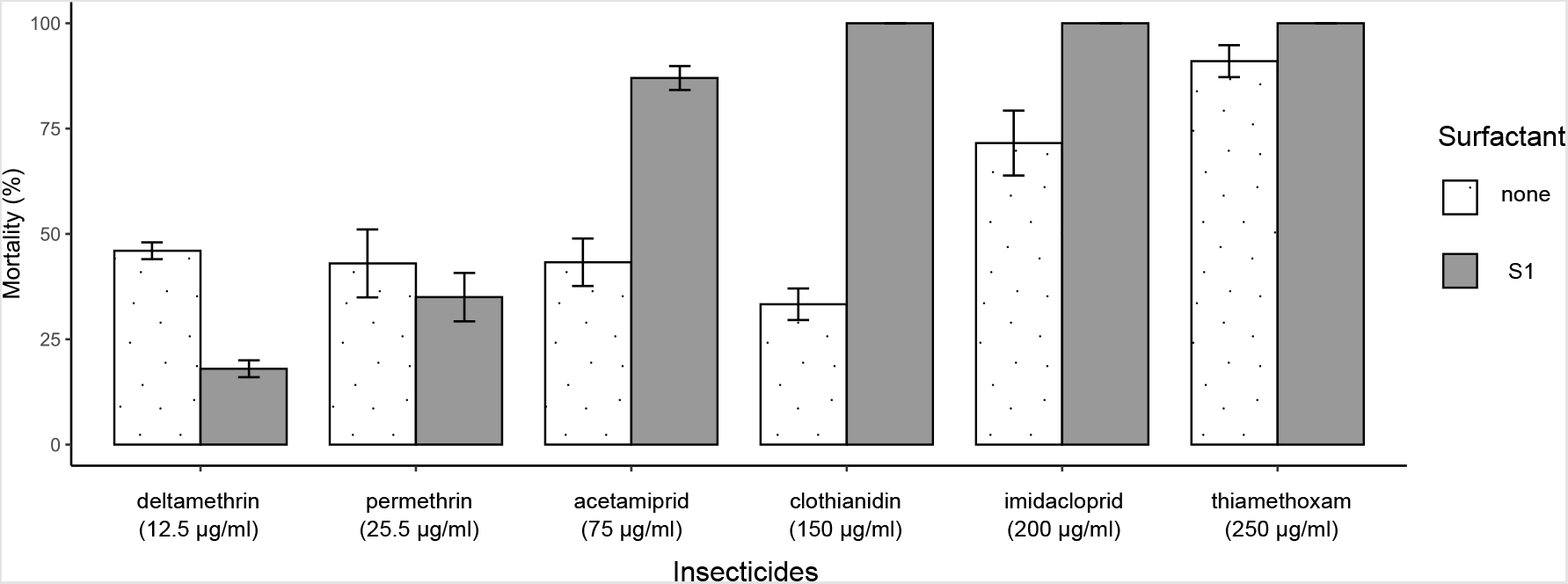
24-h mortality rates (with standard errors of the mean) induced in a multi-resistant *An. gambiae* population by six different active ingredients alone or in combination with 1% linseed oil soap. The discriminating doses are shown in brackets.

## 4. Discussion

There is a constant need to upgrade malaria vector control programs with new chemicals in order to mitigate the effects of mosquito adaptation to insecticides [7,30]. Some formulations of agrochemicals have revealed very promising results in field trials and could help reduce the negative impacts of insecticide resistance [9,31–33].

In the current study, we have shown that linseed oil soap increases the susceptibility of *An. gambiae* mosquitoes to neonicotinoids including clothianidin whose formulations have been recently approved for indoor residual spraying [7–9]. Our control experiments where female adults were exposed to linseed oil soap alone allowed us to rule out the possibility that this surfactant on its own had lethal effects on mosquitoes. Corbel et al. [14] have also demonstrated Mero®, a formulation of rapeseed fatty acid esters had no insecticidal activity on *Anopheles* mosquitoes. Using estimates of 24-h LC_50_ and LC_99_, we then found that blends of clothianidin and linseed oil soap are highly potent leading to a drastic increase in short-term lethal toxicity of the active ingredient. According to LC_50_, the combined effects of clothianidin and soap are more than 10-fold higher than that of clothianidin alone. The lethal concentrations determined by our study are in the range of values obtained when dose-response curves of *An. gambiae* mosquitoes were established using clothianidin mixed with 1% Mero® [2].

Due to synergistic interactions, the mixture between linseed oil soap and clothianidin restored neonicotinoid susceptibility in a multi-resistant *An. gambiae* population [18–20]. Similarly, a recent study has found that 100% mortality was achieved in bioassays where adult *An. gambiae* mosquitoes collected from Nkolondom were exposed to a mixture containing 150 μg/ml of clothianidin and 1% Mero® [13]. This finding suggested that other vegetable oil surfactants such as rapeseed methyl esters could also enhance the potency of clothianidin. Some organosilicon surfactants have been used in agricultural pest management to modify the surface tension of plant cells and insect cuticles in order to increase the penetration of neonicotinoid insecticides [34]. Our study and previous findings [2,12,13,30] indicate that vegetable oil esters and soap enhance the overall activity of neonicotinoids and other pesticides likely by increasing the solubility of the active ingredient and its uptake by adult mosquitoes. Indeed, we observed that linseed oil soap synergised other neonicotinoid insecticides as susceptibility to thiamethoxam, imidacloprid and to a lesser extend acetamiprid was restored with surfactant-insecticide mixtures in the multi-resistant population. In two previous studies, it has been shown that the cytochrome inhibitor piperonyl butoxide (PBO) restored susceptibility fully and partially to acetamiprid and clothianidin respectively, but had no effect on thiamethoxam [18,19]. Comparatively, vegetable oil surfactants are thus more effective than the standard synergist PBO in enhancing the efficacy of neonicotinoids.

Significant research has provided examples of synergistic interactions between adjuvants and neonicotinoids. Notably, some agricultural spray adjuvants have been shown to increase the toxicity of acetamiprid against honeybees [35]. Alkyl phenoxy polyethylene ethanol use as adjuvant in thiamethoxam formulations against the whitefly *Bemisia tabaci* resulted in 25-fold increase in the potency of the mixture compared to that of the active ingredient alone [36]. Another investigation indicated that a vegetable oil adjuvant had synergistic effects resulting in increased efficacy of thiamethoxam against cowpea thrips [37]. More generally, a review of research comparing insecticide formulations versus the active ingredient concluded that the presence of adjuvants in the formulations, in most cases, results in increased toxicity of the product formulations versus their active ingredients [15]. Overall, there is sufficient evidence, including findings from our study, that adjuvants such as surfactants can inflate the activity of chemicals leading to an overestimation of their actual efficacy against insects. This overestimation may cause bias in estimating and interpreting insecticide susceptibility in wild populations.

The standard operating procedure for testing the susceptibility of adult mosquitoes recommends using Mero® as surfactant when exposing *Anopheles* and *Aedes* populations to neonicotinoids and butenolides [4,14]. A large-scale study conducted to test the efficacy of broflanilide, a new meta-diamide insecticide, on several *Anopheles* laboratory colonies from different African countries has noted a drastic increase in toxicity of the active ingredient due to the addition of Mero® [30]. It is therefore vital to determine the insecticides or classes of insecticides for which vegetable oil surfactants may not be used in testing procedures due to potential synergistic interactions. In this study, we have tested if soap could synergize pyrethroids that are chemically very different from neonicotinoids. We observed that soap did not significantly affect the efficacy of deltamethrin and permethrin. As opposed to the synergistic effects of some surfactants on neonicotinoids, which have been relatively well studied, the interactions between vegetable oil surfactants and pyrethroids remain largely unexplored. The difference observed in this study between neonicotinoids and pyrethroids is probably due to the chemical properties of each class of insecticides. Neonicotinoids are highly water soluble compared to pyrethroids that are only slightly soluble in aqueous solutions [6]. However, in comparison to pyrethroids, neonicotinoids probably have lower binding affinity to organic materials, and as a result, when their surface tension is reduced with a surfactant, their uptake and potency increase [15]. Previous studies and our findings clearly indicate that the activity of an insecticide can be largely overestimated when the active ingredient is mixed with a surfactant. In the case of neonicotinoids, this can lead to bias estimates of susceptibility in wild insect populations.

Standard tests conducted on wild larval and adult populations showed that the activity of some neonicotinoids was below the threshold required for these chemicals to be used for vector control [1,12,18–20,28]. Most active ingredients in this class of insecticides have substantially higher lethal concentration compared with other public health insecticides [1,11,12,19,28]. In addition, bioassays have revealed reduced susceptibility to acetamiprid, thiamethoxam, imidacloprid and clothianidin in some wild populations of *An. gambiae* and *An. coluzzii* raising concern about the potential efficacy of neonicotinoids against malaria vectors [10,12,18–20]. Our results show that beside PBO, there are cheap and available synergists such as soap that could be used to enhance the potency of neonicotinoids and their potential efficacy in malaria vector control.

However, the use of synergists in susceptibility testing creates confounding effects that could negatively affect resistance management efforts. Baseline susceptibility to any active ingredient is expected to vary to some extent between wild populations and even between different laboratory colonies of a mosquito species [2,30]. The purpose of susceptibility monitoring is to gain sufficient information on the level of variability that should be taken into account in order to guide insecticide resistance management programs whose aim is to preserve the efficacy of chemical interventions [27]. For a new insecticide such as clothianidin, it is important to use sensitive tests capable of detecting early stages of resistance when effective management efforts can successfully mitigate its negative impacts.. This objective could be difficult to achieve with standard operating procedures that use extremely potent blends of the active ingredient and a synergist.

## Author Contributions

FA: Conceptualization, Formal analysis, Investigation, Methodology, Writing – original draft; CF: Conceptualization, Formal analysis, Investigation, Methodology; MA: Investigation, Methodology; VP-B, Resources, Supervision. CK: Conceptualization, Formal analysis, Funding acquisition, Investigation, Project administration, Writing – review & editing.

## Informed Consent Statement

Not applicable.

## Data Availability Statement

The data for this study have been presented within this article.

